# AlloPool: An Adaptive Graph Neural Network for Dynamic Allosteric Network Prediction in Protein Systems

**DOI:** 10.1101/2024.11.01.621466

**Authors:** M. Marfoglia, L. Guirardel, P. Barth

## Abstract

Allosteric communication is essential to protein function, facilitating the dynamic regulation of biological responses through the propagation of structural and dynamic changes between regulatory and effector sites in response to stimuli. Traditional approaches to studying protein allostery often rely on static protein structures or abstract representations involving fully connected interaction graphs, which do not capture the temporal and state-dependent nature of these dynamic systems. Here, we introduce AlloPool, a graph neural network (GNN)-based model that iteratively prunes residue interactions to identify minimal, time-dependent interaction networks that govern long-range structural and dynamic responses to chemical or mechanical stimuli. Using temporal attention and graph aggregation, AlloPool accounts for evolving protein conformations in both molecular dynamics (MD) and steered MD (SMD) simulations to predict MD and SMD trajectories. We validate AlloPool on the Pin-1 protein and the ADGRG1 and B1AR receptors, showcasing its ability to accurately recapitulate protein motions, infer allosteric communication pathways, and identify critical allosteric sites. Additionally, AlloPool identifies force-dependent changes in GAIN domain structure and reconstructs directed information flow under mechanical load. Comparative analyses indicate that AlloPool outperforms existing models in MD and SMD trajectory reconstruction, presenting a new framework for analyzing force- and ligand-induced allosteric motions. This work advances the modeling of allosteric systems and offers broad potential for applications in drug discovery, synthetic biology, and protein engineering.

## Main

Protein allostery is a fundamental regulatory mechanism in cellular biology, allowing proteins to dynamically respond to external cues like ligand binding or mechanical forces^1–3^. Traditionally, allosteric proteins have been viewed as molecular switches that shift between distinct conformational states in response to stimuli (e.g. ligand binding)^4,5^. However, advances in molecular dynamics (MD) simulations reveal a more nuanced view, uncovering complex, state-dependent allosteric behaviors that reflect the dynamic nature of protein structure and function^5–8^. These simulations show dynamically coupled allosteric residues through correlated atomic motions, providing insights into how amino acid sequence variations can modulate protein functions even at distant sites. Yet, the intricate and time-dependent nature of these interactions poses challenges for in-depth analysis. Deepening our understanding of these allosteric networks and their responses to stimuli not only advances our knowledge of protein function but also opens promising avenues potential for drug discovery, synthetic biology, and bioengineering.^4,9,10^

Computational methods are increasingly used to model allosteric networks, often leveraging graph-based frameworks that represent residues as nodes and their interactions as edges. When combined with MD simulation data, these techniques are effective for identifying long-range interactions between residues and uncovering the shortest communication paths between allosteric sites^11^. Leading methods based on Perturbation response scanning (PRS) and Mutual Information (MI) calculations utilize covariance matrices and correlation metrics to infer allosteric pathways^12–14^. While these methods provide valuable insights, they are limited by key assumptions: (i) PRS assumes linear relationships between perturbations and dynamics, failing to capture non-linear behaviors, (ii) MI analysis involves complex post-processing, which may bias the identification of the true underlying allosteric networks, and (iii) neither approach includes causal inference, relying solely on correlations without addressing directional information.

Recent advances in machine learning, particularly in deep learning and graph neural networks (GNNs), have paved the way for modeling complex, non-linear relationships and their temporal evolution. Deep learning approaches widely used in fields like weather forecasting and neuroscience, excel at capturing intricate temporal dynamics crucial for predictive accuracy^15–18^. GNNs, which operate on graph-structured data, have shown great potential for modeling dynamic interactions across domains such as social networks, traffic systems, and biological systems. In particular, neural relational inference (NRI) learns interpretable embeddings that capture interaction patterns from motion data, making it useful for identifying allosteric pathways in MD simulations by uncovering latent residue interactions^19^. However, NRI assumes a static interaction graph, limiting its capacity to account for time-dependent conformational changes, particularly in non-equilibrium simulations like Steered MD (SMD). To address this shortcoming, a dynamic NRI (dNRI) was developed, allowing for the learning of multiple interaction graphs over time and offering improved performance^15^. Nonetheless, both NRI and dNRI rely on fully connected graphs, which fail to explicitly account for structural changes that alter physical interactions between neighboring residues.

In this study, we introduce *AlloPool*, a GNN-based model that refines protein interaction networks by iteratively pruning edges to capture essential residues involved in dynamic and allosteric processes. Using temporal attention mechanisms and graph aggregation layers, *AlloPool* adapts its interaction graph to reflect changes in protein conformation over time. Validation across MD and SMD simulations—including for the Pin-1 protein, the ADGRG1 GAIN domain, and the GPCR beta 1 adrenergic receptor (B1AR)— demonstrates *AlloPool*’s ability to capture dynamic shifts in allosteric networks and predict mutation impacts. Additionally, *AlloPool* facilitates unsupervised classification of protein states by detecting interaction pattern changes, establishing it as a valuable tool for investigating allostery and designing targeted, state-specific interventions in protein systems.

### Overview of function, architecture, and rationale

Traditional machine learning models for studying protein allostery or force propagation pathways have typically focused on one specific mechanism, often failing to generalize between ligand-induced allosteric communication and force-induced mechanical signaling. While allosteric models concentrate on how information transfer and conformational changes impact protein function, force propagation models investigate how mechanical forces are transmitted within a protein structure. Despite the apparent distinction between these mechanisms, we hypothesized that both rely on the same fundamental principle: the transmission of dynamic changes across long-range networks. This shared principle suggests the possibility of a unified framework capable of capturing both allosteric and force propagation pathways.

To address this, we propose *AlloPool*, a graph neural network (GNN)-based model that bridges the gap between ligand-induced and force-induced communication mechanisms by learning minimal and dynamic interaction graphs that evolve over time. Unlike previous methods that assume static interaction networks, *AlloPool* iteratively prunes non-essential edges, focusing on the smallest subset of interactions necessary to explain protein dynamics across different conformational states. By incorporating temporal attention and graph aggregation layers, the model is able to learn time-dependent interaction graphs that reflect the structural and functional changes induced by external forces or ligand binding.

The core architecture of *AlloPool* is designed to learn transition interaction graphs that capture the minimal set of edges driving protein dynamics (Fig. 1). The model processes time series data from MD and SMD simulations, transforming each frame into a graph where nodes represent α-carbons and edges represent spatial interactions between residues. These graphs are input into an encoder-decoder framework, where a temporal attention layer first condenses the time series into a compact representation. The encoder then applies successive edge pruning layers, removing edges that do not contribute to the system’s dynamics. This pruning results in a minimal interaction graph, which is passed to the decoder for trajectory reconstruction. Through this process, *AlloPool* identifies the key changes in interaction networks associated with shifts in allosteric communication or force propagation.

**Figure 1.**
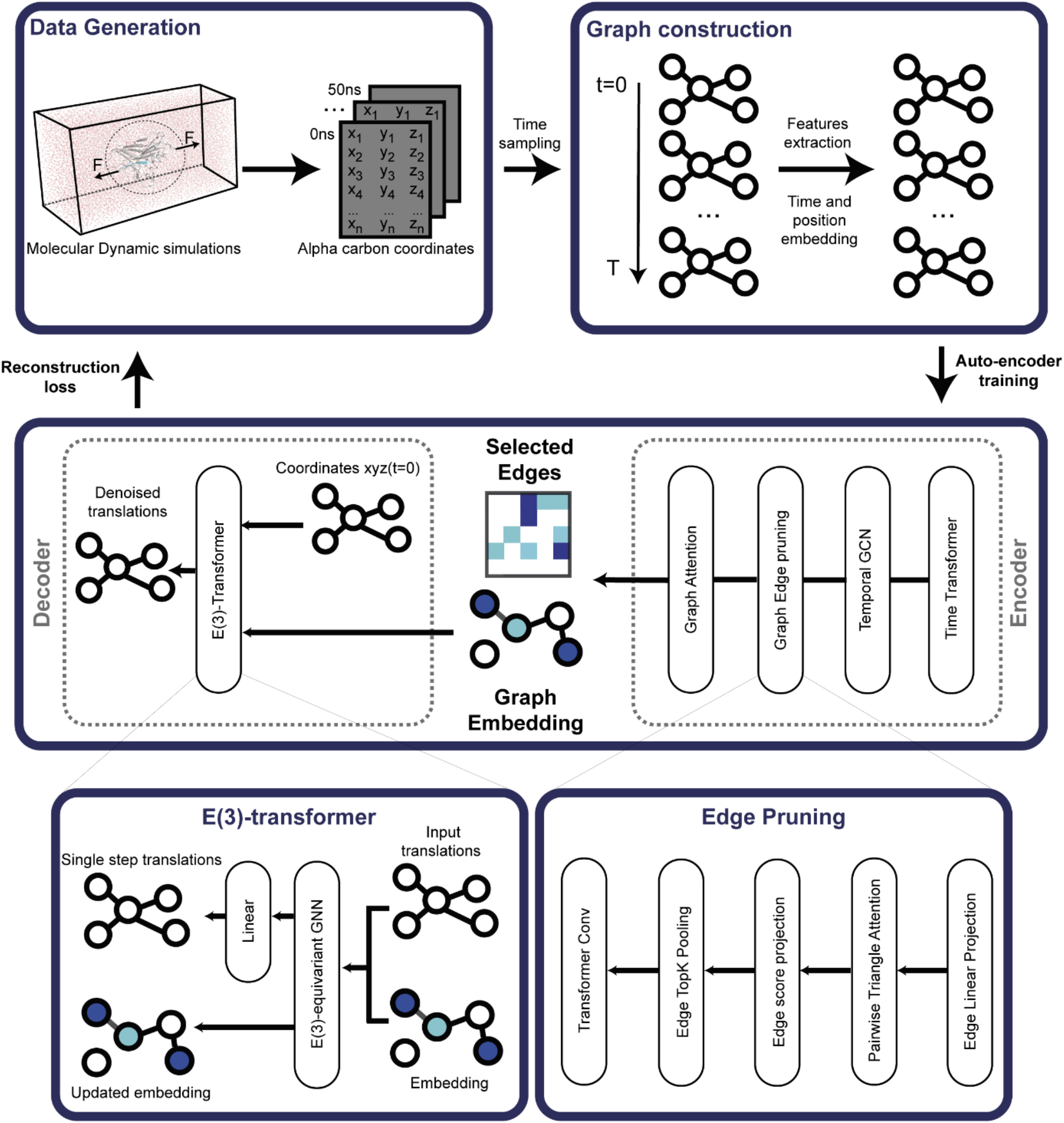
AlloPool learns allosteric interaction networks from dynamic data. Detailed architecture of the AlloPool network. Simulations are performed to obtain dynamic trajectories of a system (data generation). Each time point of the trajectory is represented by a graph with associated node and edge features with an adjacency matrix. Frames are sampled from the trajectory and used to train the AlloPool model with two jointly trained components. First, an encoder is trained to select relevant edges from the adjacency matrix generating the latent embedding that depicts the allosteric network of the system. Meanwhile, a decoder is trained to predict the future translations and reconstruct the observed trajectories of the system given the latent representation generated by the encoder. The allosteric process is thus described as the minimal set of learned edges that enable accurate prediction of the system’s dynamic compared to the true trajectories.

Principal component analysis (PCA) is applied to the learned interaction graphs to detect shifts in dynamic modes over the simulation trajectories, revealing significant changes in residue interactions that correspond to conformational shifts. This enables the identification of critical regions within the protein structure that are involved in allosteric signaling or mechanical force transmission. By reconstructing force propagation pathways, *AlloPool* can map out how mechanical forces are transmitted through the protein under load, demonstrating its capability to capture both allosteric and mechanical signaling mechanisms in a unified framework.

### AlloPool learns force propagation pathways in the GAIN domain

ADGRG1, an adhesion G-protein coupled receptor (aGPCR), provides a valuable test case for evaluating AlloPool due to its dual role in ligand-induced allosteric signaling and force-induced mechanosensing^20–22^. ADGRG1 mediates critical processes in neuronal development and cancer, responding to chemical and mechanical stimuli, making it ideal for testing AlloPool’s adaptability. Prior research shows ADGRG1 undergoes ligand-binding conformational shifts that initiate signaling events, as well as mechanical force propagation through defined pathways. Using MD and SMD data, we evaluate AlloPool’s capacity to capture and interpret these complex information flows within ADGRG1’s mechanosensitive GAIN domain, whose isolated structure has been experimentally characterized.

To evaluate AlloPool’s capability in identifying critical allosteric connections (edges) in non-equilibrium simulations, we conducted Steered Molecular Dynamics (SMD) simulations on the GAIN domain of ADGRG1 by applying forces at both N- and C-termini with pulling speeds of 0.1 Å/ns and 1 Å/ns (Fig. 2A, Methods). The resulting simulation trajectories were used to train AlloPool as per the architecture in Fig. 1. Upon pulling, mechanical stress accumulated within the GAIN domain (Fig. 2B), leading to conformational changes and partial unfolding of its first helix (Fig. 2E-F), which formed new inter-helix contacts at the N-terminus of GAIN^21^. Correlation-based force propagation pathways were computed as reference, allowing us to compare these with AlloPool’s edge selections and confirm the model’s accuracy in capturing relevant dynamic allosteric connections.

**Figure 2.**
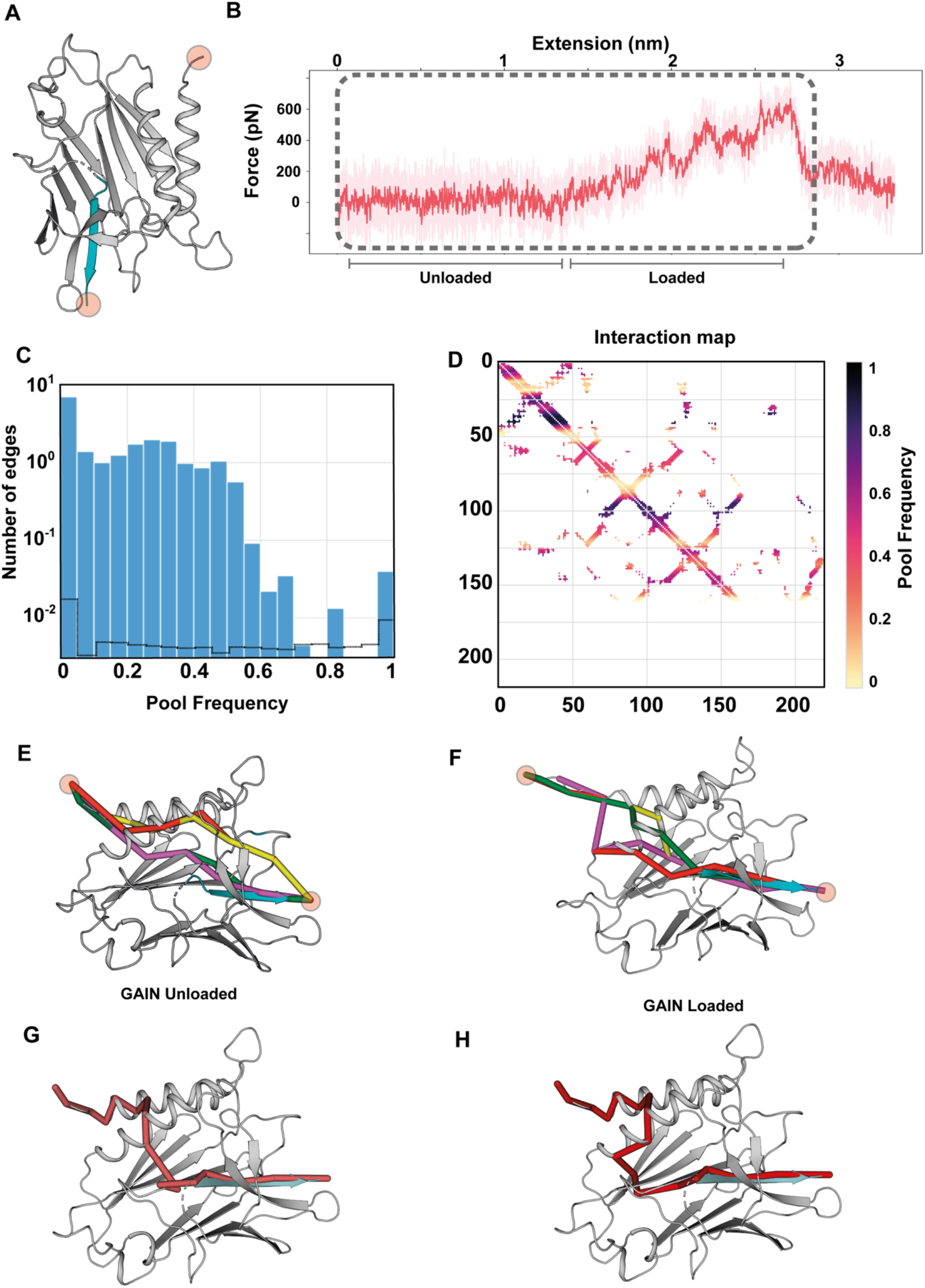
AlloPool learns allosteric interaction networks enabling force propagation pathway identification. (A) Structure of the GAIN domain with the 2 residues onto which the pulling force is applied highlighted by red spheres. (B) Representative SMD trajectory (dark red) and rolling average (light red) leading to the loaded state. The region of the SMD trajectory used to train the model is represented by a gray dotted box. Unloaded and loaded states’ region are depicted by gray lines. (C) Distribution of the pooling frequencies represented as histogram compared to the edge persistence distribution (gray dotted line). (D) Allosteric interaction network output by the model. The x-y axes represent the residue index. (E-F) Best-scoring force propagation pathways reconstructed from the interaction map for the unloaded (left) and loaded (right) state respectively. (G-H) Force propagation pathways for unloaded (left) and loaded (right) states obtained from classic correlation analysis.

Training the model on early segments of the simulations (prior to the initial unfolding event) enabled it to reconstruct trajectories with high fidelity, achieving an average RMSD accuracy of 1.3 Å, even in the absence of explicit structural inputs. We examined the frequency of edge selections by AlloPool and compared this to the persistence of residue interactions throughout the simulations (Fig. 2C). Persistent interactions showed a binary distribution, either absent or present, while edge selection frequencies by AlloPool followed a 1-inflated binomial distribution (Fig. 2C), suggesting that the model prioritized edges that are critical to the protein’s dynamic states. High-frequency edges aligned with contiguous backbone connections linking adjacent residues but also captured critical tertiary contacts between distal residues, even without direct sidechain information (Fig. 2D). Importantly, the interaction network map (Fig. 2D) showed asymmetry, illustrating the model’s capacity to create directed edges, thus reflecting a directional flow of allosteric information across the structure.

To evaluate the functional relevance of the selected edges (Fig. 2D) in response to applied force, we reconstructed potential force propagation pathways between the N- and C-termini of the GAIN domain (Methods). Using the rationale described previously^21^, we extracted these pathways for both unloaded and loaded phases of the simulations, prioritizing routes with high average pool frequencies along each path. Remarkably, pathways connecting the residues at the force application sites aligned closely with those identified through correlation-based methods (Fig. 2E-H, indicating that AlloPool effectively identifies edges critical for modulating the GAIN domain’s dynamics under mechanical stress. Additionally, the observed shift in pathway structure between unloaded and loaded states reveals that AlloPool is accounting for structural changes induced by force. These findings suggest that AlloPool successfully identifies force propagation pathways in the GAIN domain, capturing physically-meaningful interactions even when structural alterations occur due to mechanical load.

### Low-dimensional decomposition of selected edges identifies shift in the system’s dynamics

While the trained model successfully learned GAIN conformational dynamics from the SMD simulations, we further investigated whether dynamic mode shifts could be identified directly from the learned edges and if they reflected any intrinsic molecular switching properties of the system. We performed PCA on both persistent and pooled edges (Fig. 3A, B) extracted from the trajectory ensemble and decomposed the pooled edge time series using non-negative matrix factorization (NMF). The PCA projection of persistent edges defined distinct simulation trajectories (Fig. 3A), whereas the pooled edges correlated with time, indicating that the pooling strategy and learned edges primarily reflect the intrinsic dynamics of the GAIN domain rather than variations in SMD trajectories (Fig. 3B).

**Figure 3.**
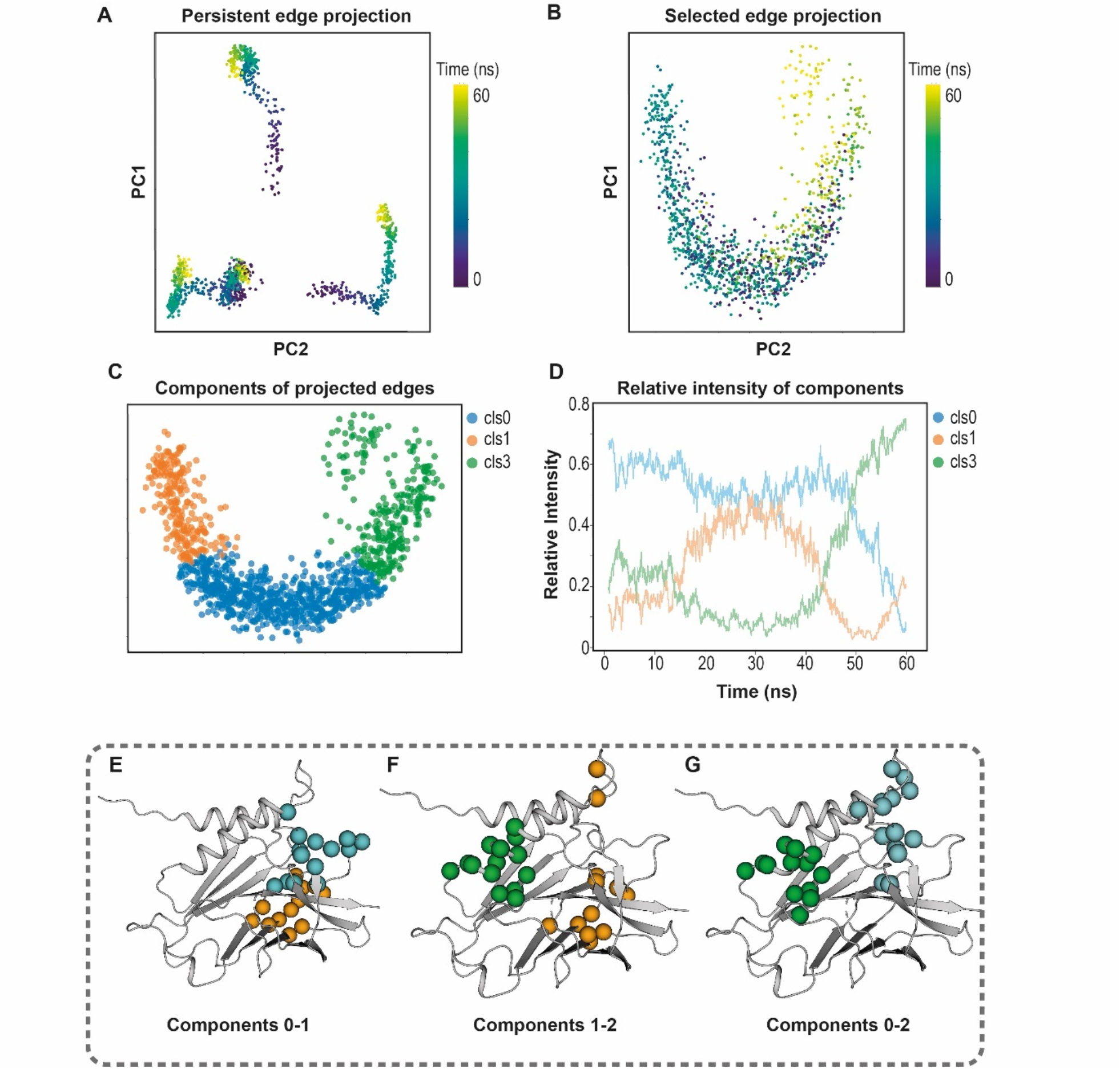
Selected edges identify force-dependent shifts of dynamics. (A-B) PCA of persistent edges (A) or selected edges (B) represented over the two principal components and colored by time. (c) Results of the non-negative matrix factorization clustering of the selected edges representing the three identified clusters. (d) Evolution of the relative intensity of the NFM components over time. (E-G) Comparison of the directed edges that significantly contribute to the discrimination across components 0 and 1 (E, blue: C0-C1, orange:C1-C0), components 1 and 2 (F, orange: C1-C2, green: C2-C1) and components 0 and 2 (G, blue: C0-C2, green: C2-C0).

To evaluate dynamic shifts in edge selection, we decomposed the projected edges using NMF (Fig. 3C) and analyzed their relative intensities over time (Fig. 3D). The components revealed three major dynamic ensembles, each with fluctuating intensities throughout the trajectory ensemble. Component 1 (blue) remains relatively stable until it decreases around 50 ns, while Component 2 (orange) peaks between 20-35 ns, and Component 3 (green) dominates towards the end of the simulation, after 35-40 ns.

To understand which edges influenced this component distinction, we extracted the most discriminatory edges and mapped the involved nodes onto the GAIN structure (Fig. 3E-G). Comparing Components 1 and 2, we found that Component 1 (blue spheres) differentiated itself through key edges linking helices H1 and H2, as well as loops 7-10 of the GAIN domain (Fig. 3E). In contrast, Component 2 (orange spheres) highlighted connections between β-strands 3-4 and 9, stabilized by disulfide bridges within the GPS motif that enhance the thermodynamic stability of the GAIN domain. Meanwhile, nodes differentiating Component 3 (Fig. 3F-G) were predominantly connections between helix H2 and β-strands 5-6, reflecting polar interactions stabilizing the α-domain helices with β-strand 6 (Fig. 3G).

These findings demonstrate that the model not only captured critical allosteric networks from the simulation trajectories but also identified distinct dynamic modes that depend on different types of interactions, effectively tracking their evolution under mechanical load.

### AlloPool learns key allosteric residues in Pin-1 and outperforms existing methods

To evaluate AlloPool’s performance relative to models utilizing fully connected graphs, we compared AlloPool with the Neural Relational Inference (NRI) model on the allosteric peptidyl-prolyl cis/trans isomerase Pin-1 using simulation data from previous publication^19^. Three Pin-1 states were selected for the comparison: the apo form (PDB: 3TDB), an agonist-bound state (PDB: 1NMV), and an antagonist-bound state (PDB: 1NMV). Both AlloPool and NRI models were independently trained on each Pin-1 state with parameters defined for each respective model (Methods). Additionally, NRI was trained on GAIN domain trajectories to benchmark its performance against AlloPool for interaction graph learning in non-equilibrium simulations.

To compare the models’ abilities to reconstruct protein dynamics, we calculated the standard deviation of the reconstructed trajectories’ Root Mean Square Fluctuation (RMSF) values. To confirm that each model accurately recapitulates the structural dynamics of Pin-1, we also computed the Root Mean Square Deviation (RMSD) between the true and reconstructed trajectories. The results for average RMSD and RMSF are presented in Table 1.

**Table 1.** Comparison of AlloPool and NRI performance for reconstructing trajectories. Comparison of average root mean squared deviation (RMSD, Å) of alpha carbons and value standard deviation (VSD, Å) of root mean squared fluctuations (RMSF). The best model value for these two metrics is shown. NRI: classic NRI default parameters. AlloPool parameters : conditions of time sampling (number of frames + interval between them) defined in this study.

| **System** | **Pin-1 Apo** | **Pin-1-FFpSPR** | **Pin-1-pCdc25C** | **GAIN** |
| --- | --- | --- | --- | --- |
| **NRI RMSD** | 4.83 Å | 1.63 Å | 5.72 Å | **1.43 Å** |
| **NRI SVD** | 0.39 Å | 0.24 Å | 0.9 Å | 0.16 Å |
| **AlloPool parameters-NRI RMSD** | 2.74 Å | 1.83 Å | 5.93 Å | 1.69 Å |
| **AlloPool parameters-NRI SVD** | 0.12 Å | 0.11 Å | 0.06 Å | 0.05 Å |
| **AlloPool RMSD** | **1.49 Å** | **1.02 Å** | **3.07 Å** | 1.59 Å |
| **AlloPool SVD** | **0.11 Å** | **0.11 Å** | **0.04 Å** | **0.05 Å** |

Overall, our AlloPool models outperformed the NRI models across these systems. While NRI models were able to replicate trajectory fluctuations as measured by RMSF, we observed a notable variability in RMSD errors, ranging from 1.4 Å to 5.72 Å across different NRI models. This variability suggests that NRI models may capture intrinsic system fluctuations but fail to accurately reconstruct the translational displacements observed between frames in the simulation. Here, we define translations as the displacement in Cartesian space between two simulation frames.

To investigate the discrepancy between RMSD and RMSF further, we calculated the norm of the predicted translations (Fig. 4A-B). Remarkably, AlloPool trained on Pin1-Apo predicted translations that aligned closely with the observed values (within 1 Å), while NRI-predicted translations were three times higher than AlloPool’s denoised translations and 4.5 times higher than the true observed values. This finding indicates that AlloPool provides more realistic translations that maintain the protein fold structure.

**Figure 4.**
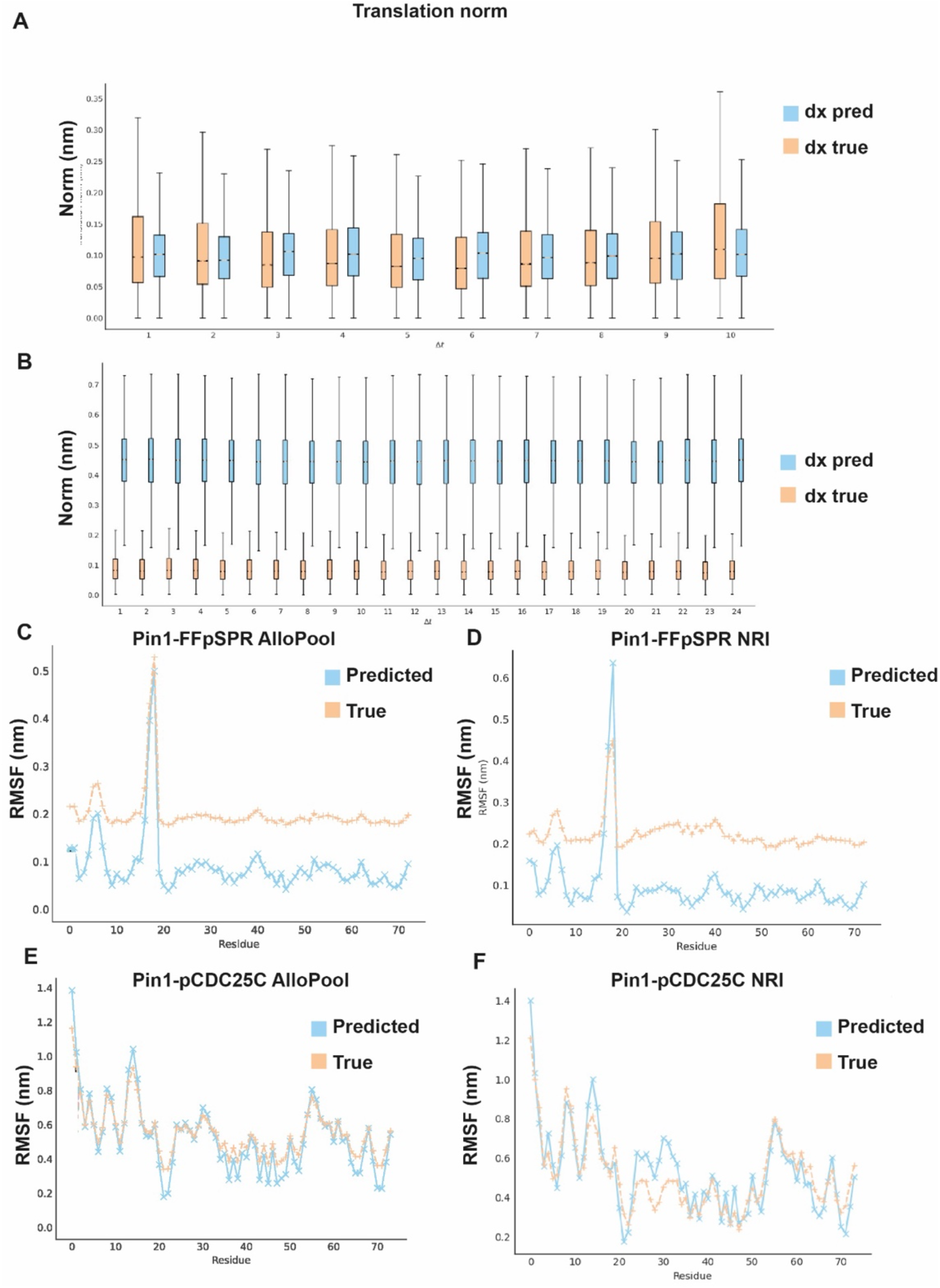
AlloPool outperforms the NRI in predicting translations and dynamics. **(A-B)** Norm of the translations denoised by AlloPool (A) or NRI (B) models. (C-F) RMSF plots for the reconstructed and true trajectories for AlloPool (C-E) and NRI (D-F) models for Pin1-FFpSPR and Pin1-pCDC25C.

**Figure 5.**
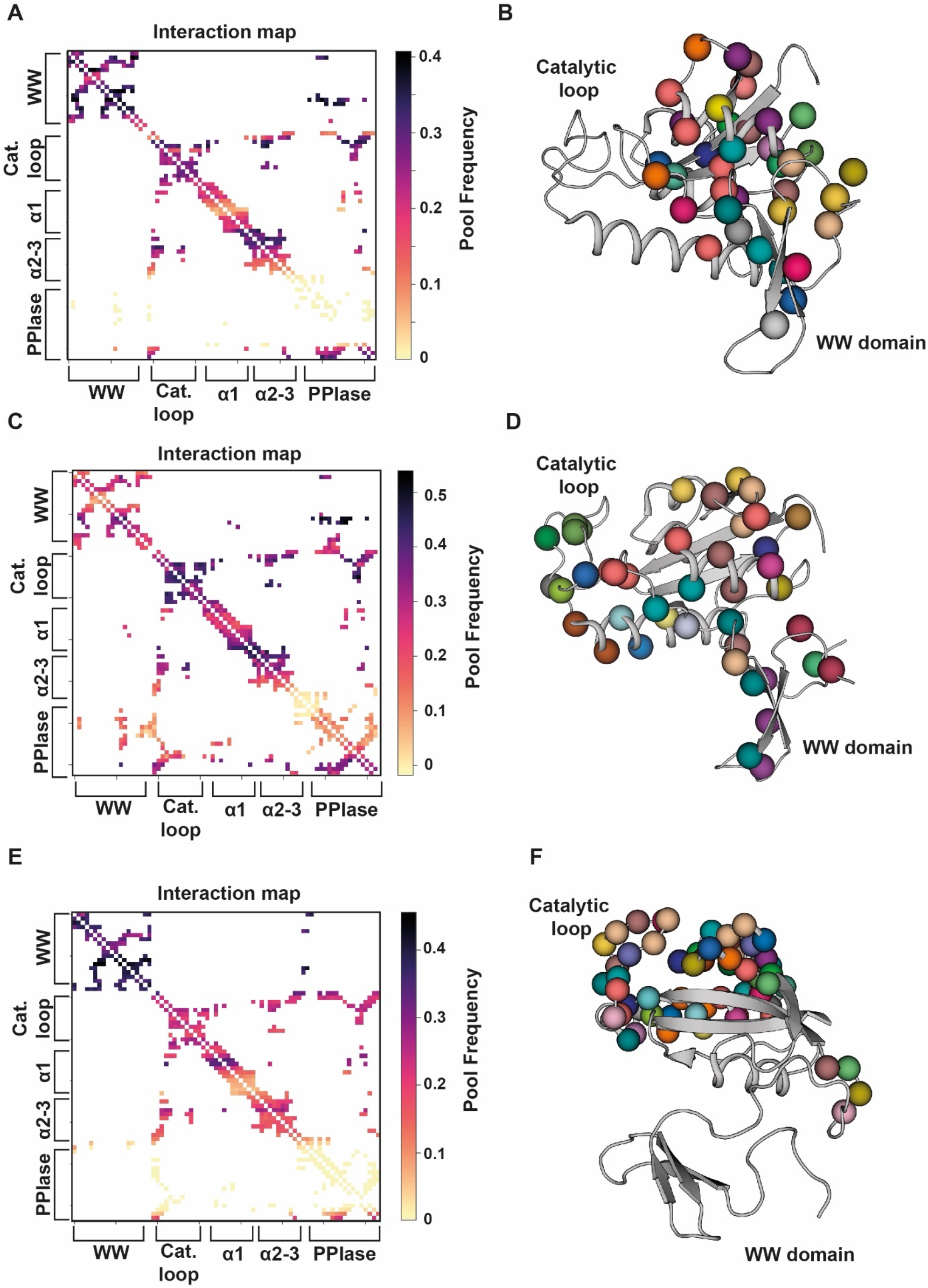
AlloPool recapitulates interdomain allostery in Pin1. (A,C,E) Interaction maps learned for Pin-11 Apo (A), agonist-bound (C) and antagonist-bound (E) trajectories. (B,D,F) Best allosteric residues pairs found for Pin-1 apostate (B), agonist-bound (D) or antagonist-bound (F) are represented by spheres.

We then compared the reconstructed RMSF values with those from the NRI model (Fig. 4C-F). AlloPool more accurately recapitulated native fluctuations within the trajectory while utilizing fewer edges in its predictions. Additionally, the NRI model tended to exaggerate fluctuations in the loop regions of Pin-1, a phenomenon not observed in AlloPool’s predictions (Fig. 4C-F).

### AlloPool learns allosteric sites in GPCRs

To further assess AlloPool’s ability to capture complex allosteric coupling, we benchmarked its performance on G-protein-coupled receptors (GPCRs), a class of allosteric proteins that switch between distinct conformational-functional states upon ligand binding. Structural studies of GPCRs have identified conserved residue “microswitches” that undergo conformational changes between states, such as the DRY, PIF, and W-toggle motifs—key sites driving the transition between inactive and active receptor states in class A GPCRs. In this study, we focused on the beta-1 adrenergic receptor (B1AR) and simulated six unique B1AR-ligand pairs, each representing distinct functional states, to evaluate AlloPool’s ability to predict allosteric coupling across these states. For each ligand-bound B1AR simulation, AlloPool was trained on individual trajectories, after which the learned allosteric interaction graphs were analyzed.

AlloPool, when trained on B1AR-agonist trajectories, accurately identified interactions involving the canonical R.3.50 (DRY motif), P5.50 (PIF motif), and the W-toggle motif, connecting them with extracellular ligand and intracellular G-protein binding surfaces (Fig. 6A). This result demonstrates AlloPool’s ability to capture established GPCR allosteric signatures with high fidelity, underscoring its applicability to proteins with well-defined allosteric sites.

**Figure 6.**
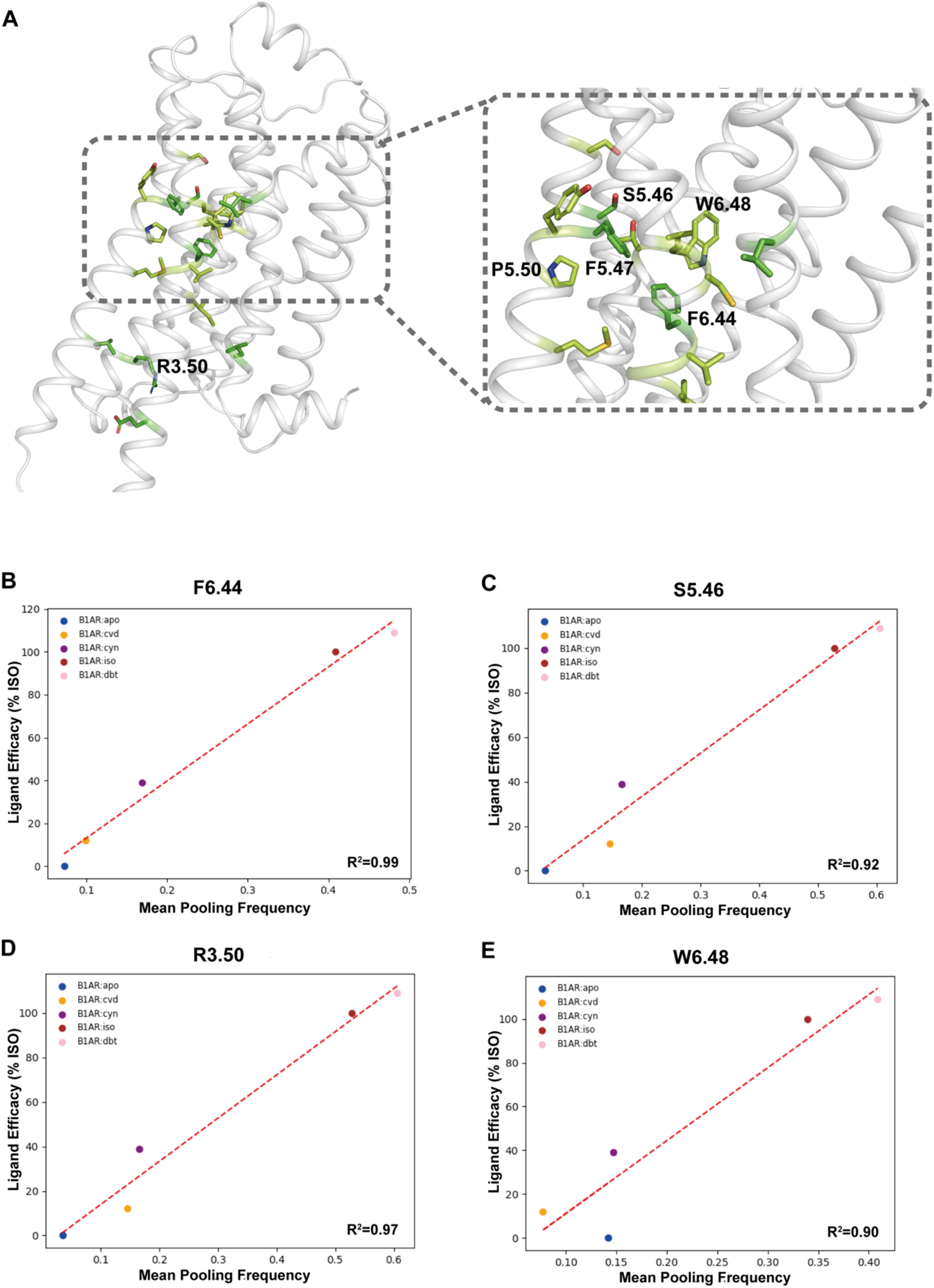
Prediction of allosteric sites in the B1AR receptor. (A) Key allosteric interactions identified by AlloPool, colored by averaged pooling frequency (light green: > 50%, green: >75%) for the agonist-bound B1AR. (B-C) Linear regression analysis of ligand efficacy (y-axis) as a function of the average pooling frequency. Data points are colored by B1AR:ligand pairs. The red dashed line represents the linear regression fit to the data, along with the associated R^2^ score.

We next explored AlloPool’s capacity to differentiate structural and dynamic perturbations caused by ligands with varying efficacies (i.e., a ligand’s propensity to activate specific receptor functions). Our analysis revealed a strong correlation between ligand efficacy profiles and distinct allosteric interactions identified by the model (Fig. 6B-C), indicating that AlloPool can pinpoint critical sites that mediate receptor activation while capturing ligand-induced structural and dynamic changes. Among these interactions, AlloPool identified residues in direct ligand contact, suggesting that B1AR’s dynamic response inherently reflects ligand chemical properties—despite the model’s lack of explicit ligand structural information during training. This finding highlights AlloPool’s ability to infer ligand-specific interactions through dynamic residue behavior alone (Fig. 6C, S5.46), highlighting its potential for broadly predicting functionally relevant allosteric networks in ligand-receptor systems.

## Discussion

The AlloPool model establishes a new standard in modeling time-dependent protein allosteric behaviors by dynamically identifying minimal interaction networks necessary for accurate trajectory prediction, a significant improvement over static, fully connected graph models. This enables AlloPool to adaptively capture chemically- and mechanically-induced allosteric effects within a unified framework. Findings from simulations of Pin-1 and ADGRG1 highlight AlloPool’s ability to account for conformational shifts and non-equilibrium states, providing valuable insights into the temporal nature of allosteric communication.

AlloPool’s temporal attention mechanism and graph aggregation layers enable an evolving representation of protein interactions that aligns with structural rearrangements in response to both ligand binding and mechanical forces. Its capacity to model both chemical and mechanical allostery is evidenced in simulations of adhesion protein ADGRG1 and ligand-binding protein B1AR, supporting an integrative approach to understanding allosteric mechanisms across diverse protein families and broadening its applications in drug design for both ligand-sensitive and mechanosensory targets.

Notably, AlloPool’s high precision in mapping force propagation pathways under Steered MD (SMD) simulations on the GAIN domain illustrates its effectiveness in capturing residue-residue interactions that mediate force transmission, a critical feature in mechanosensory proteins. The directed, asymmetric interaction networks observed further suggest AlloPool’s ability to infer directional flow in response to force, excelling in dynamic contexts where traditional correlation-based methods often fall short. This promising result underscores AlloPool’s potential to predict transient, non-equilibrium states as well as equilibrium behaviors.

Comparative analyses with NRI models underscore AlloPool’s superior performance in both root mean squared deviation (RMSD) and root mean squared fluctuation (RMSF) metrics, with reduced network complexity achieved through selective edge pruning. This targeted approach enhances dynamic prediction accuracy, especially in systems with high structural variability. Additionally, non-negative matrix factorization (NMF) on selected edges reveals distinct dynamic modes that align with force-dependent structural shifts, validating AlloPool’s ability to capture temporally resolved allosteric dynamics.

The model’s performance across different Pin-1 states points to promising applications in studying proteins with multiple functional conformations, presenting opportunities to identify state-specific allosteric sites. AlloPool’s potential for unsupervised state classification through interaction network changes introduces a powerful tool for designing interventions that modulate protein function, with significant implications in drug design and synthetic biology.

AlloPool effectively addresses core limitations in allosteric modeling by utilizing a dynamic, edge-pruning GNN architecture that captures the intrinsic time-dependent nature of protein function. Future directions include expanding AlloPool’s application to diverse protein families to assess its performance in wider functional contexts and exploring its integration with experimental data to enhance predictive accuracy. Additionally, incorporating advanced causal inference techniques could further refine its ability to capture directional interactions in protein networks. By advancing our understanding of allosteric and force-dependent mechanisms, AlloPool presents a robust approach to modeling complex biological systems, with significant implications for both basic research and therapeutic development.

## Author contributions

PB and MM conceived the project. MM and LG developed the model. MM, LG and PB analyzed data. PB and MM wrote the manuscript.

## Methods/ Supplementary

### Molecular Dynamics simulations

The data used to train and validate the model were obtained from Molecular Dynamics (MD) simulation at 20 ns per step.

Our models are trained on simulation windows of ten frames, regularly spaced every ten frames. Each window therefore covers a simulation time of 20 × 10 × 9 = 1800 ns between its first and last frame.

The model is evaluated on a simulation on which it has not been trained.

### Feature processing

Features consist in the spatial coordinates of the residue C*α*, spatial coordinates of the residue C*β* (or H*α*3 in the case of glycine), as well as sinus and cosine of backbone dihedral angles. We also add the C *α* displacement between consecutive frames, which can serve as an approximation for atomic velocity. There are thus 13 node features, to which we add a 64-dimensional absolute positional embedding that represents the residue index in the sequence and directionality of the amino.

The input therefore consists of a window of *T* = 10 frames with *N* nodes represented in dimension *d* = 77. The training set contains *B* windows depending on the length of the MD or SMD simulations.

We then standardize the data feature-wise to 0 mean and 1 standard deviation.

We denote 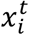 the set of features of residue *i* at frame 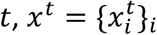 the set of all residue features at frame *t*, and 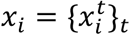 the trajectory of residue *i*. We also denote 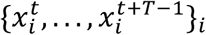 the time window starting at time *t*.

### Persistence of edges

Our model processes frame data at each time window *t* as a graph 𝒢_*t*_(*V, E*_*t*_), with *N* the set of residues. Because a complete graph would be too computationally expensive to process and would not make physical sense (no physical contacts between selected edges), we chose to define edges based on residue distances. More precisely, (*i, j*) ∈ *E*_*t*_ if *d*(*i, j*) < 1 nm for at least 30% of the frames in the time window [*t, t* + *T* − 1].

### Edge Pool model

Our edge pool model, following classic architecture of autoencoders, simultaneously learns edges of the protein graph driving its dynamics, and the dynamics of that protein, in an unsupervised manner.

This model minimizes the RMSD between *x*^*t*^ and 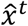 for *t* ≥ 1. We compute the RMSD using the Kabsch-Umeyama algorithm to align the structures prior to loss computation. We formalize it as 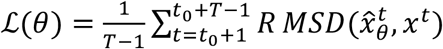 where θ are the trainable parameters of the model *h*.

The model is composed of an encoder module, and a E(N)-Transformer decoder module.

The encoder itself is composed of three sub-modules: the time processing module, the edge pruning module, and the graph processing module.

The time processing module consists of four Transformer layers, followed by Temporal Graph Convolution Layers. These layers are applied independently to each node and learn a temporal weight for each node independently, that is used to reduce the time dimension from *T* frames to one by taking the product of node representations multiplied by nodes’ temporal weights. The output of the time processing module is therefore *N* nodes with dimension *d*_1_.

Then, this output is passed to the edge pooling module consisting of 6 successive Edge-TopK blocks. Each of these blocks passes the input graph through two SageConv layers, which update node representations, then compute scores for each edge using a learned weight *w* of dimension 2*d*_1_, using this procedure:

Let 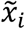 be the output features of the SageConv layers for node *i*, of dimension *d*_1_. Let *E*_*t*_ be the edges associated to this graph. Then, the Edge-TopK block operation is as follow:

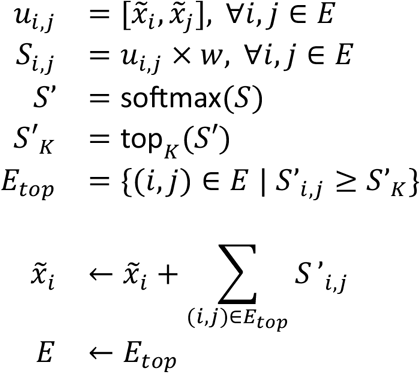

To introduce some randomness and avoid the model getting stuck, nodes features are dropped randomly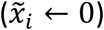 to zero before each scoring, with a probability p=0.01.

After the pooling module, the resulting graph is passed to the structural processing module, composed of 1 Graph Convolution layer, using the edges conserved by the pooling layer, to perform message passing.

We thus obtain an intermediate representation *x*^′^, with *N* nodes with features of dimension *d*_2_.

This intermediate representation is then fed to the E(N)-Transformer of depth T-1 which will predict translations from x(t=0) to x(t+1), from x(t+1) to x(t+2), and so on, until the final translation x(t) to x(t+T).

The model weights are optimized using the Adam optimizer with an initial learning rate of 10^−4^, a weight decay coefficient of 0.1, and a learning rate scheduler decreasing the learning rate by 2 after 100 optimizer steps with no loss decrease. The windows are grouped in batches of 10.

### Training situations

In the following, *g*_θ_ represents the encoder and *h*_θ_ the decoder of the model with their respective parameters.

The encoder takes as input a simulation window 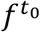 and outputs a single representation *g*_θ_, while the decoder outputs a reconstituted time window 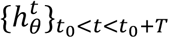.

### Trajectory prediction objective

In the first training situation, the model learns to reconstitute the trajectory. At each training time window 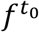 and for each time step *t*, we predict:

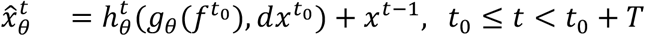

where 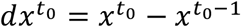.

### Trajectory denoising objective

In a second training situation, the model learns to denoise the trajectory. At each training time window 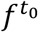 and for each time step *t*, we predict:

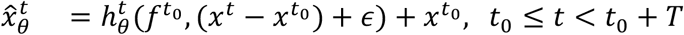

where ϵ ∼ 𝒩(0,1)

### Edge factorization

To analyze trends in selected edges across the trajectories, we use Non-negative Matrix Factorization (NMF) to decompose the edge selection matrix 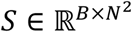 defined as:

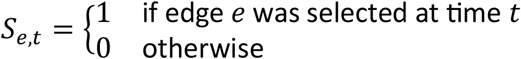

NMF decomposes *S* as such: *S* ≃ *WH*, where 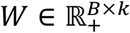 and 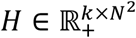 have non-negative coefficients.

We used the NMF implementation from the scikit-learn library, with *α*_W_ = 0, *alpha*_*H*_ = 10^−>^, *l*1_*ratio*_ = 0.8.

By multiplying *H* column-wise by 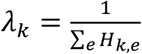 and *W* row-wise by 1/λ_*k*_, we rescale *H* so that its columns sum to 1, without modifying the decomposition *WH* ≃ *S*. This way, the coefficients *h*_*k,e*_ represent the importance of edge *e* in component *k*. We can attribute an edge *e* to its “main component” *k*_*e*_ = *argmax*_*k*_ *h*_*k,e*_.

### Pathway calculation

First, we can assign each edge a global importance score with its edge selection probability 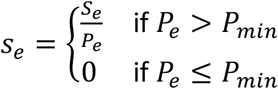 where *S*_*e*_ (resp. *P*_*e*_) is the number of training windows where edge *e* has been selected (resp. is a persistent edge), and *P*_*min*_ = *B*/10 is defined as 10% of the total numbers of training windows.

Then, we convert edge scores to edge weights 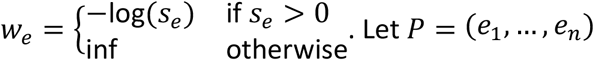 be a path of length *n*. We define path score 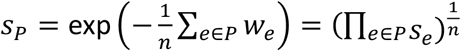.

Finally, we take the shortest paths according to edge weights *w*_*e*_ between to sets of residues corresponding to source and target protein regions, and sort them according to path scores.

